# Node abnormality predicts seizure outcome and relates to long-term relapse after epilepsy surgery

**DOI:** 10.1101/747725

**Authors:** Nishant Sinha, Yujiang Wang, Nádia Moreira da Silva, Anna Miserocchi, Andrew W. McEvoy, Jane de Tisi, Sjoerd B. Vos, Gavin P. Winston, John S. Duncan, Peter Neal Taylor

**Author notes:** (NS); (PNT).

## Abstract

**Objective:** We assessed pre-operative structural brain networks and clinical characteristics of patients with drug resistant temporal lobe epilepsy (TLE) to identify correlates of post-surgical seizure outcome at 1 year and seizure relapses up to 5 years.

**Methods:** We retrospectively examined data from 51 TLE patients who underwent anterior temporal lobe resection (ATLR) and 29 healthy controls. For each patient, using the pre-operative structural, diffusion, and post-operative structural MRI, we generated two networks: ‘pre-surgery’ network and ‘surgically-spared’ network. The pre-surgery network is the whole-brain network before surgery and the surgically-spared network is a subnetwork of the pre-surgery network which is expected to remain unaffected by surgery and hence present post-operatively. Standardising these networks with respect to controls, we determined the number of abnormal nodes before surgery and expected to remain after surgery. We incorporated these 2 abnormality measures and 13 commonly acquired clinical data from each patient in a robust machine learning framework to estimate patient-specific chances of seizures persisting after surgery.

**Results:** Patients with more abnormal nodes had lower chance of seizure freedom at 1 year and even if seizure free at 1 year, were more likely to relapse within five years. In the surgically-spared networks of poor outcome patients, the number of abnormal nodes was greater and their locations more widespread than in good outcome patients. We achieved 0.84 ± 0.06 AUC and 0.89 ± 0.09 specificity in detecting unsuccessful seizure outcomes at 1-year. Moreover, the model-predicted likelihood of seizure relapse was significantly correlated with the grade of surgical outcome at year-one and associated with relapses up-to five years post-surgery.

**Conclusion:** Node abnormality offers a personalised non-invasive marker, that can be combined with clinical data, to better estimate the chances of seizure freedom at 1 year, and subsequent relapse up to 5 years after ATLR.

## Introduction

Epilepsy surgery is an effective treatment for bringing seizure remission in drug-resistant focal epilepsies, however, it is underutilised (Wiebe *et al.*, 2001; Langfitt and Wiebe, 2008; Vakharia *et al.*, 2018). One reason for the under-referral of patients is the reservations regarding the uncertainty of its outcome (Haneef *et al.*, 2010; Vakharia *et al.*, 2018). In around 30-40% of cases, seizures continue despite surgery and the multidisciplinary team are unable to accurately predict this risk prior to surgery (Janszky *et al.*, 2005; Spencer and Huh, 2008; Téllez-Zenteno and Wiebe, 2008; de Tisi *et al.*, 2011; Bell *et al.*, 2017). Therefore, to better inform this clinical decision making, there is a need to predict seizure outcomes both in the short-term, and the likelihood of seizure relapse in the long term (Cohen-Gadol et al., 2008; de Tisi et al., 2011).

The incomplete removal of a wider epileptogenic network is increasingly being recognised as one of the reasons for continued seizures post-surgery (Spencer, 2002; Richardson, 2012). Many studies, driven by the aforementioned hypothesis, have attempted predicting seizure outcomes from pre-surgical data (Bonilha *et al.*, 2015; Munsell *et al.*, 2015; Goodfellow *et al.*, 2016; Bell *et al.*, 2017; Keller *et al.*, 2017; Morgan *et al.*, 2017; Proix *et al.*, 2017; Sinha *et al.*, 2017; Gleichgerrcht *et al.*, 2018). Most studies, however, have investigated brain networks without incorporating the knowledge of the planned/performed surgery into the analysis. Naturally, the outcome of epilepsy surgery will depend not only on the pre-surgery brain network, but also on how the surgery (i.e., its location and extent) will affect the brain network (Taylor *et al.*, 2018). Including surgical data allows the inference of a ‘surgically-spared’ network – the subnetwork for which none of the connections are altered by surgery and are therefore expected to remain after the surgery. Thus, the presence of epileptogenic structures in the surgically-spared network, a likely cause for seizure relapse in short or long term after surgery, needs investigation.

Studies employing quantitative imaging have consistently demonstrated that in temporal lobe epilepsy there are structural abnormalities that involve brain structures beyond the hippocampus and the temporal lobe (Bernasconi *et al.*, 2004; Keller and Roberts, 2008; McDonald *et al.*, 2008; Concha *et al.*, 2012; Otte *et al.*, 2012; Deleo *et al.*, 2017). Accumulating evidence suggests that these abnormalities configure a network of abnormal structures that may be involved in the generation of seizures (Spencer, 2002; Bonilha *et al.*, 2013; Bernhardt *et al.*, 2015; Bonilha and Keller, 2015). Indeed, the pathophysiological mechanisms associated with epileptogenesis have a strong basis in aberrant neural connectivity (Liu *et al.*, 2016; Besson *et al.*, 2017). Therefore, quantifying the abnormalities before, and expected to remain after, surgery may inform postoperative seizure outcome.

The main goal of our study was to understand how structural network abnormality related to seizure outcomes after temporal lobe epilepsy surgery. We investigated the abnormality of the ‘surgically-spared’ networks because, at a conceptual level, post-operative outcomes will likely be determined by what remains post-surgery. Our study addresses three main questions: a) do patients with more abnormalities have worse postoperative seizure outcomes? b) does surgery have a greater effect on node abnormality in seizure-free patients? c) if the node abnormality measure is to be used alongside common clinical variables of a patient, would it generalise to make patient-specific predictions on the chances of seizure freedom after surgery? Our study shows that the node abnormality is an important measure to be considered alongside other pre-surgical clinical factors to evaluate the risk of poorer seizure outcomes in patients with refractory TLE.

## Methods

### Participants

We studied 51 patients who underwent unilateral anterior temporal lobe resection at the National Hospital of Neurology and Neurosurgery, London, United Kingdom and 29 healthy controls. Presurgical clinical information included: sex, age at epilepsy onset, age at surgery, epilepsy duration, number of AEDs taken before surgery, history of status epilepticus, history of secondary generalised seizures, side of surgery, evidence of MRI pathology, evidence of hippocampal sclerosis, history of depression, history of psychosis, and history of any other psychiatric disorders. Patients were followed up after surgery and classified according to the ILAE scale of seizure outcome at 12-month intervals (Wieser *et al.*, 2001). One year after the surgery, 34 patients were completely seizure free (ILAE 1), 8 patients continued to have auras only (ILAE 2), and 9 patients were not seizure free (ILAE 3-6). This group of patients constitute the same cohort as in our previous study (Taylor *et al.*, 2018).

ILAE surgical outcomes of seizure freedom were recorded annually at years 1 and 2 for all 51 patients, at year 3 for 45 patients, year 4 for 37 patients, and at year 5 for 31 patients. We considered that a patient had a seizure relapse if, at any given year after year 1, the ILAE outcome of the patient changed from ILAE 1-2 to ILAE 3-6. If the ILAE outcome of a patient did not change to ILAE 3-6 and the follow-up duration was less than five years, it cannot be ascertained if the patient would have relapsed upon a full five-year follow-up. Therefore, beyond the known follow-up period, we did not include such patients in our analysis. The full preoperative clinical information, postoperative ILAE outcomes, and relapse data for each patient is provided in Table S1.

We used *χ*^2^ test to check for the differences between the outcome groups in sex, side of surgery, evidence of hippocampal sclerosis, evidence of MRI pathology, history of status epilepticus, evidence of generalised seizures, history of depression, psychosis, and any other psychiatric disorders. Differences in age at epilepsy onset, age at surgery, epilepsy duration, and number of anti-epileptic drugs taken preoperatively between the outcome groups, were assessed with two-tailed non-parametric Wilcoxon ranksum test. Table 1 shows the summary of patient details.

**Table 1:** Demographic and clinical data of patients

| *Year 1 outcome*<br>*Variables* | *ILAE 1* | *ILAE 2* | *ILAE 3-6* | *Significance* |
| --- | --- | --- | --- | --- |
| *Patients (n)* | 34 | 8 | 9 |  |
| *Sex*<br>*(Male/Female)* | 16/18 | 2/6 | 2/7 | $\chi^2_{1,2} = 0.54, p_{1,2} = 0.46$<br>$\chi^2_{1,3+} = 0.93, p_{1,3+} = 0.33$<br>$\chi^2_{2,3+} = 0.19, p_{2,3+} = 0.66$ |
| *Age at Onset*<br>*(mean ± std)* | 12.2 ± 10.3 | 14.2 ± 11.4 | 19 ± 12 | $p_{1,2} = 0.62$<br>**$p_{1,3+} = 0.04$**<br>$p_{2,3+} = 0.43$ |
| *Age at Surgery*<br>*(mean ± std)* | 38 ± 11.9 | 38.6 ± 10.3 | 46.5 ± 10.2 | $p_{1,2} = 0.96$<br>$p_{1,3+} = 0.08$<br>$p_{2,3+} = 0.13$ |
| *Epilepsy Duration*<br>*(mean ± std)* | 25.8 ± 15.8 | 24.3 ± 17.3 | 27.5 ± 7.3 | $p_{1,2} = 0.74$<br>$p_{1,3+} = 0.54$<br>$p_{2,3+} = 0.54$ |
| *Side*<br>*(Left/Right)* | 22/12 | 3/5 | 5/4 | $\chi^2_{1,2} = 1.02, p_{1,2} = 0.31$<br>$\chi^2_{1,3+} = 0.01, p_{1,3+} = 0.91$<br>$\chi^2_{2,3+} = 0.06, p_{2,3+} = 0.79$ |
| *Hippocampal sclerosis* | 24 (70.5%) | 6 (75%) | 5 (55.5%) | $\chi^2_{1,2} = 0.03, p_{1,2} = 0.85$<br>$\chi^2_{1,3+} = 0.21, p_{1,3+} = 0.64$<br>$\chi^2_{2,3+} = 0.11, p_{2,3+} = 0.74$ |
| *Number of AEDs before surgery*<br>*(mean ± std)* | 6.3 ± 2.4 | 7.6 ± 2.9 | 9.2 ± 3.3 | $p_{1,2} = 0.10$<br>**$p_{1,3+} = 0.01$**<br>$p_{2,3+} = 0.46$ |
| *Preoperative MRI*<br>*(Normal/Abnormal)* | 5/29 | 1/7 | 2/7 | $\chi^2_{1,2} = 0.16, p_{1,2} = 0.69$<br>$\chi^2_{1,3+} = 0.001, p_{1,3+} = 0.97$<br>$\chi^2_{2,3+} = 0.01, p_{2,3+} = 0.91$ |
| *History of status epilepticus* | 5 (15.7%) | 0 (0%) | 3 (33.3%) | $\chi^2_{1,2} = 0.30, p_{1,2} = 0.58$<br>$\chi^2_{1,3+} = 0.63, p_{1,3+} = 0.43$<br>$\chi^2_{2,3+} = 1.35, p_{2,3+} = 0.25$ |
| *Secondary generalised seizures* | 28 (82.3%) | 6 (75%) | 6 (66.7%) | $\chi^2_{1,2} = 0.0005, p_{1,2} = 0.98$<br>$\chi^2_{1,3+} = 0.32, p_{1,3+} = 0.57$<br>$\chi^2_{2,3+} = 0.02, p_{2,3+} = 0.88$ |
| *Depression* | 8 | 5 | 2 | $\chi^2_{1,2} = 2.96, p_{1,2} = 0.09$ |
| | | | | $\chi^2_{1,3+} = 0.13, p_{1,3+} = 0.72$<br>$\chi^2_{2,3+} = 1.42, p_{2,3+} = 0.23$ |
| *Psychosis* | 2 | 0 | 0 | $\chi^2_{1,2} = 0.05, p_{1,2} = 0.83$<br>$\chi^2_{1,3+} = 0.02, p_{1,3+} = 0.88$ |
| *Other psychiatric disorders* | 8 | 3 | 4 | $\chi^2_{1,2} = 0.13, p_{1,2} = 0.72$<br>$\chi^2_{1,3+} = 0.68, p_{1,3+} = 0.41$<br>$\chi^2_{2,3+} = 0.04, p_{2,3+} = 0.84$ |
| *Relapsed by end of Year 2* | 0 (0%) | 3 (37.5%) | 9 (100%) |  |
| *Relapsed by end of Year 3* | 3 (8.8%) | 5 (62.5%) | 9 (100%) |  |
| *Relapsed by end of Year 4* | 5 (14.7%) | 6 (75%) | 9 (100%) |  |
| *Relapsed by end of Year 5* | 7 (20.6%) | 6 (75%) | 9 (100%) |  |

As a control group, we studied 29 healthy individuals, with no significant medical history of neurological or psychiatric problems. The control group was age and gender matched to the patient group. Control data was used as a normative measure for the metrics obtained from patients. The study was approved by the NHNN and the Institute of Neurology Joint Research Ethics Committee, and written informed consent was obtained from all subjects.

### MRI acquisition, data processing, and surgery network

For each patient in this study, T1-weighted structural (sMRI) and diffusion-weighted (dMRI) data were acquired before the surgery. Within one year after the surgery, T1-weighted sMRI data was acquired again for each patient. Detailed imaging protocols are described in Supplementary Methods. Postoperative sMRI was used to accurately delineate the resected tissue. Complete details on how we drew the resection mask for these subjects are published in (Taylor *et al.*, 2018). Briefly, the postoperative sMRI data was linearly registered to the preoperative sMRI using FSL, then the patient-specific resection masks were manually drawn, and finally validated by two independent raters for a majority subset.

Next, we applied the same data processing pipeline as in our previous study (Taylor *et al.*, 2018) to incorporate the information of surgery for inferring the two networks: whole-brain pre-surgery network and surgically-spared subnetwork. Different processing steps involved in generating these networks are detailed in Supplementary Methods. In brief, the preoperative sMRI data were parcellated into 114 cortical and subcortical regions of interest (ROIs) derived from the predefined Geodesic Information Flow atlas and separately in 82 ROIs using the Freesurfer Desikan-Killiany atlas in the native space of each participant. We registered the parcellated ROIs, resection mask, and tracts from deterministic tractography on dMRI data in native space. The pre-surgical streamline network is the connectivity matrix depicting the number of streamlines connecting two ROIs. The surgically-spared streamline network is inferred after removing the streamlines that intersected the resection mask. By definition, surgery can only cause an immediate reduction in the number of streamlines. Therefore, we specified that the surgically-spared network contains only those network edges which are not expected to change in streamline count following surgery (i.e., edges where their streamlines do not pass through/into the resection cavity). These concepts are illustrated in Figure 1.

**Figure 1:**
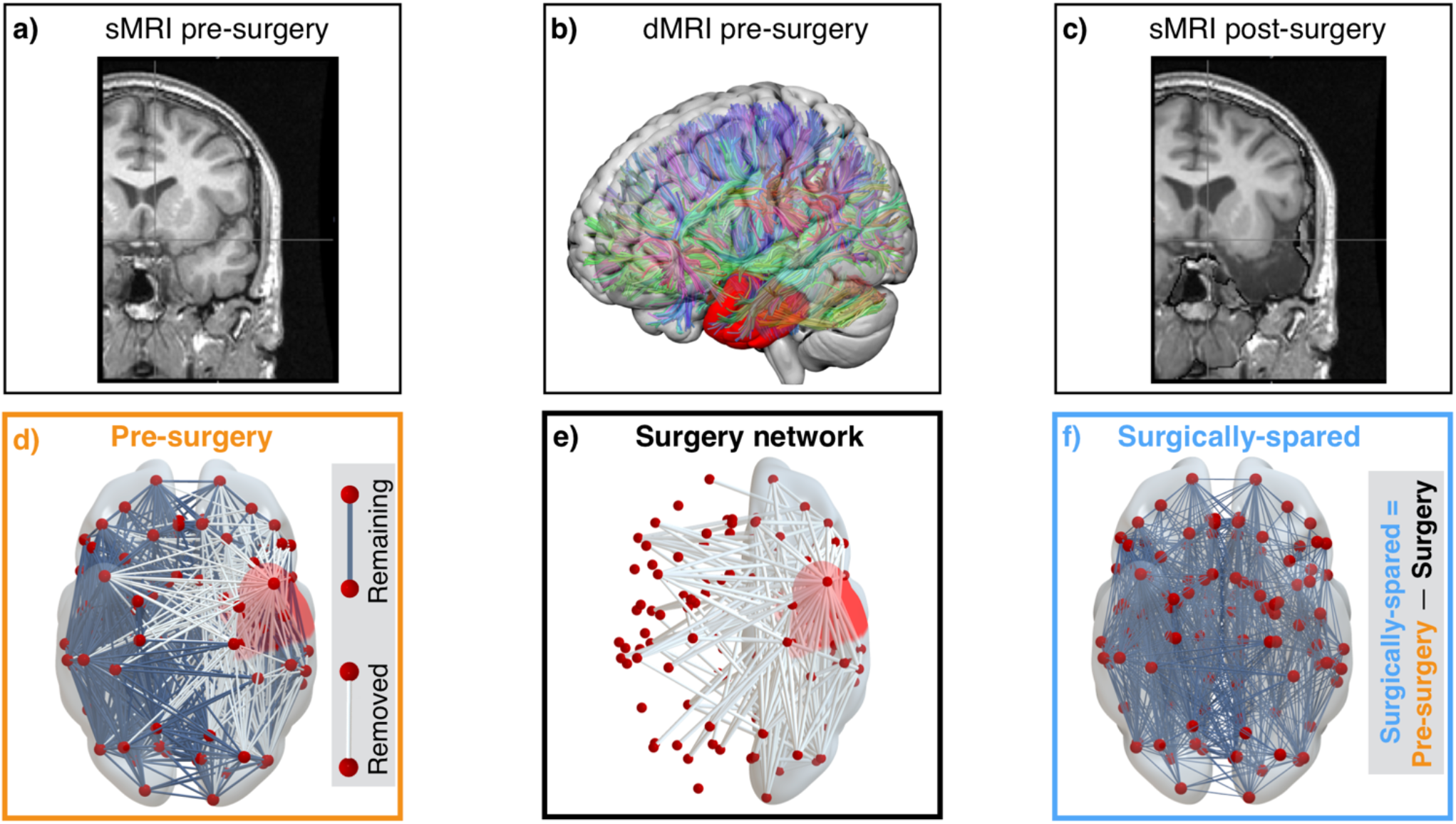
Estimating patient-specific surgery network. Preoperative T1w MRI of an example patient in panel **a)** and postoperative T1w MRI in panel **c)** were used to delineate the tissue resected by surgery. The resected tissue shown by the red resection mask in panel **b)** was used with the preoperative diffusion MRI to infer brain networks. Pre-surgery network inferred based on the number of streamlines connecting different ROIs in panel **d)** ignores the surgery information by not taking the resection mask into consideration. The patient-specific surgery network is illustrated in panel **e)** which shows the connections that were affected by the surgery. Panel **f)** shows the surgically-sparred subnetwork.

T1-weighted MRI and dMRI data were also acquired for 29 control participants using the same MRI scanner and imaging protocols as the patients.

### Node abnormality computation

The overall pipeline summarising different steps to compute node abnormality is illustrated in Figure 2. For each subject, we inferred networks based on the mean generalised fractional anisotropy (gFA) property of the streamlines obtained from the dMRI data (Tuch, 2004). We standardised the pre-surgery gFA network of each patient against controls as follows: for each connection present between ROIs *i* and *j* in a patient, the connection distribution was obtained from the equivalent connection between ROIs *i* and *j* from the control networks. The z-score for that connection was calculated as the number of standard deviations away from the mean, where the standard deviation and mean were obtained from the control distribution. Networks inferred from deterministic tractography are sparse, so we z-scored only those connections in patients for which an equivalent connection existed in at least 10 (∼35%) controls (de Reus and van den Heuvel, 2013). This is depicted in Figure 2a.

**Figure 2:**
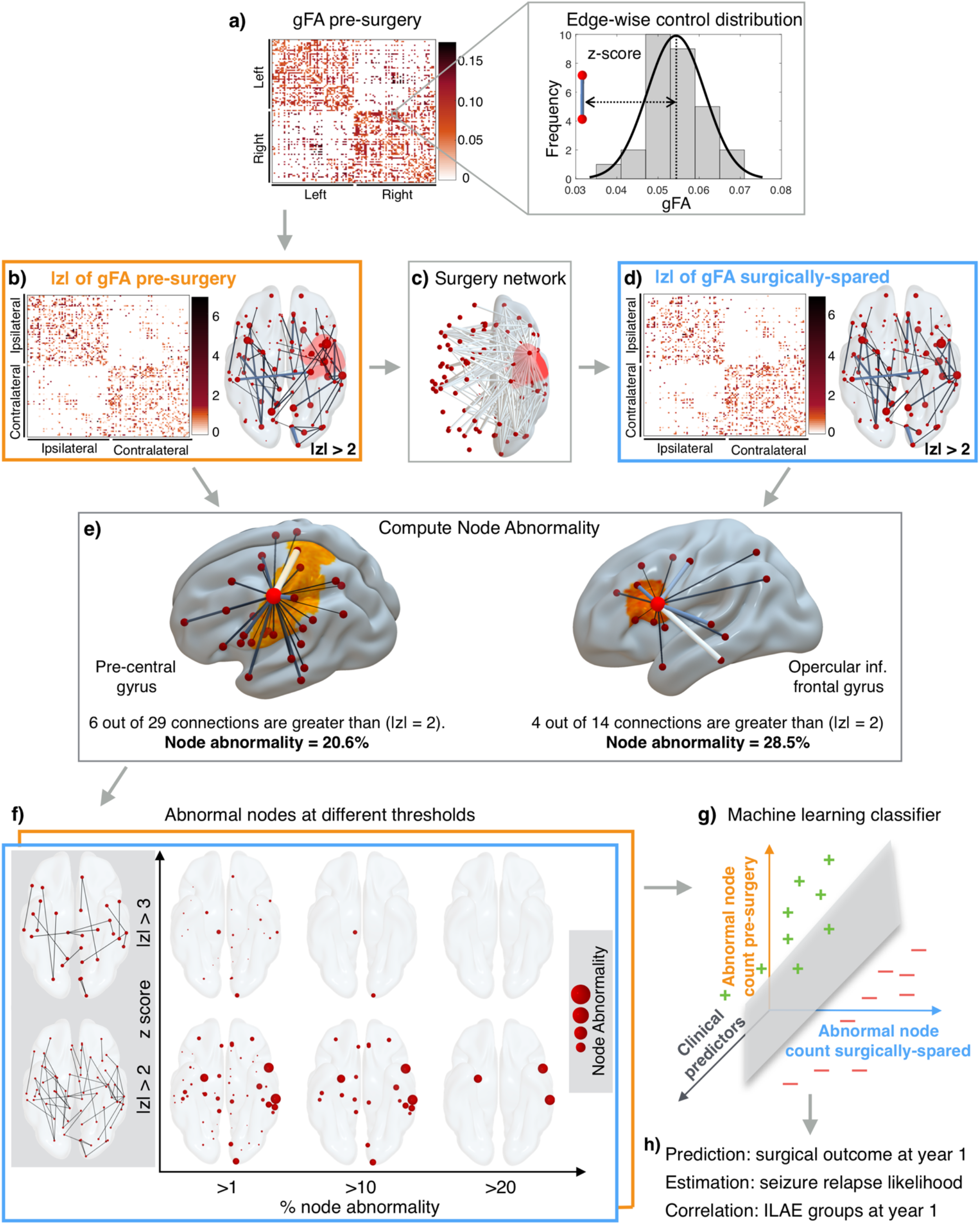
Overall pipeline. Pre-surgical gFA network architecture for each patient in panel **a)** is inferred and edges are standardised against a control distribution to obtain a z-score transformed network in panel **b)**. The connections affected by the surgery shown in surgery network in panel **c)** are removed to obtain surgically-spared network in panel **d)**. Panel **e)** shows the concept of node abnormality for two example nodes. By normalising the number of abnormal links to a node with its degree, the heterogeneity in the degree of network nodes is accounted for. A high degree node can be less abnormal compared to a low degree node depending on the number of abnormal connections. **f)** Different thresholds required for the computation of node abnormality are shown. z-score at which a link is considered abnormal is on the y-axis and the cut-off at which a node is considered abnormal is shown on the x-axis. **g-h)** Abnormal node count in pre-surgery and surgically-spared networks are incorporated in a machine learning classifier along with the clinical predictors to predict surgical outcomes with its severity at year-one and estimate chances of seizure relapse in five years.

After obtaining the pre-surgery z-scored gFA network, we removed the connections present in the surgery-affected network to obtain the surgically-spared z-transformed gFA network. High |*z*| indicates high deviation from normality. Thus, the pre-surgery network maps the abnormal links present before the surgery and surgically-spared network maps the abnormal links that would remain unaffected immediately after the surgery. This is illustrated in Figure 2b-d.

To study how different regions (nodes) are affected in these networks, we computed node abnormalities (Figure 2e) in the pre-surgery and surgically-spared networks by counting the number of abnormal links to each node. We normalised the number of abnormal links to a node by its degree in the pre-surgical network, thus expressing node abnormality in percentage terms.

Quantification of node abnormality load raises two questions: first, what is the definition of an abnormal connection? second, when is a node considered abnormal? The former is essential for the application of a threshold on the abnormality network to count the number of abnormal links at each node. For the latter, another threshold is needed to define beyond what percentage level a node should be considered abnormal. We therefore varied the z-score threshold from 2.1 to 4.5 in increments of 0.1 and the percentage abnormality threshold from 1% to 50% in increments of 1%. At each point on this two-dimensional grid, we computed how many nodes were abnormal in pre-surgical and surgically-spared networks. This is illustrated in Figure 2f for six example threshold pairs. Finally, having quantified the abnormality load for each patient, we assessed its discriminatory ability in predicting seizure outcomes and seizure relapse.

### Quantifying the change in abnormality load after ATLR

To investigate the effect of surgical treatment on reduction of abnormalities, we compared the change in abnormality load between the pre-surgery and the surgically-spared networks. The ROIs in the left and right hemispheres of patients were expressed as ipsilateral or contralateral to surgery. Then, we categorised each ROI into 6 ipsilateral and 6 contralateral areas i.e., temporal, subcortical, parietal, occipital, frontal, and cingulate cortices. In each area, we determined the number of abnormal nodes in the pre-surgery and surgically-spared networks patient-specifically. Then, we averaged the number of abnormal nodes in each area for the seizure free (ILAE 1) and non-seizure free (ILAE 3-6) groups of patients. Finally, by computing the proportion of abnormal nodes in every area (i.e., ratio of mean abnormal ROIs to the total number of ROIs in each area) for the pre-surgery and surgically-spared network, we noted the change due to the surgery in all patients.

### Predictive model design for generalizability assessment

We predicted the patient-specific probability of seizure relapse using preoperative clinical data, pre-surgery node abnormality, and the surgically-spared node abnormality. We performed this using support vector machine (SVM) implemented in MATLAB ‘fitcsvm’ classification library (Platt, 1999; Guyon *et al.*, 2002). We applied a linear kernel because this enables the interpretation of weight vectors (i.e., the relative importance of each feature in the prediction), which were used to rank the importance of metrics in identifying patients who would have suboptimal seizure outcome. SVMs were initially trained with all 15 preoperative metrics: 13 clinical, 1 pre-surgery node abnormality, and 1 surgically-spared node abnormality. To identify the most informative metrics, after each round of SVM training, we removed the least important metric (in terms of its weight vector) and trained a new SVM with the remaining metrics. We repeated this process until only a single metric remained (Guyon *et al.*, 2002; Fagerholm *et al.*, 2015). At each stepwise removal we recorded: a) the performance of classifier in classifying totally seizure free (ILAE 1) and non-seizure free (ILAE 3-6) patients, and b) the Spearman’s rank correlation between the predicted probability of seizure relapse for each patient with the actual severity of seizure outcomes at one-year after surgery (ILAE class).

The performance of the classifier was estimated using binary classification. Given that ILAE 2 patients tend to relapse (Table 1, S1), and thus, are in the spectrum between the totally seizure free (ILAE 1) and non-seizure free (ILAE 3-6) patients, we first excluded the ILAE outcome group 2 patients (Fairclough *et al.*, 2018). With these patients removed, our dataset consisted of 43 samples, 34 of which were labelled 1 corresponding to ILAE 1, and 9 were labelled −1, corresponding to ILAE 3-6. On this dataset, we performed nested-cross validation by combining a three-way split of the data (training-validation-testing) with leave-one-out cross-validation (CV) and grid search for SVM parameter (box-constraint) tuning. This was done to avoid upward bias in the metrics of performance estimates (Guyon and Elisseeff, 2003; Tsamardinos *et al.*, 2018). Additionally, we avoided any bias in the selection of the most discriminatory threshold pair (i.e., *z-*score and percentage abnormality) to determine the node abnormality by computing it at every step of cross-validation after removing the test subject (Smialowski *et al.*, 2009).

Specifically, in nested-cross validation, an external leave-one-out is implemented in which one patient is left out at every step for testing and the remaining patients used for training and validation. Training and validation were performed in the internal leave-one-out CV in which one patient is again left out for validation and the remaining used for model training combined with model parameter tuning. In our analysis, we tuned the model on 100 logarithmically spaced grid points between 1 and 10. At every point, the SVM is trained and its performance tested using the patient left out for validation by estimating AUC. We selected the model parameter that gave the highest cross-validated AUC. The classifier generalisation capability is then evaluated by computing the classification AUC, accuracy, sensitivity, and specificity using the patient originally left for testing in the external cross-validation. We also noted the probability with which each test patient was classified as non-seizure free. The intuition being that the predictive model, though blind to the non-seizure free outcome categories (i.e., all ILAE 3 to ILAE 6 are labelled as −1), would classify the patients with worse surgical outcome with a higher probability.

To determine where the ILAE outcome group 2 subjects fall on the spectrum, we treated all 8 ILAE outcome group 2 patients as test subjects. SVMs were trained and tuned, as described above, on all the remaining seizure free (ILAE 1) and non-seizure free (ILAE 3-6) patients (43 patients). On the classifier with highest discrimination between the seizure free and non-seizure free patients, we tested the features of ILAE 2 patients to note only the probability of classification to the non-seizure free group. We refer to these probabilities as the likelihood of seizure relapse because a high probability indicates a predicted propensity towards a non-seizure free outcome. Having obtained the likelihood of seizure relapse for all 51 patients, we compared this with the surgical outcome categories at year 1 and the actual seizure relapse in five years post-surgery. Note that the labels for all training data are binary and based on 12-month ILAE1 versus ILAE3-6 outcomes only. The model is therefore blind to severity of outcome (i.e. ILAE class 2, 3, 4, 5), and also blind to outcomes beyond 12 months.

### Statistical analysis and data availability

To investigate if a greater number of abnormal nodes are associated with suboptimal seizure outcomes post-surgery, we applied the non-parametric Wilcoxon rank-sum test. One-tailed *p* value was computed using the ranksum function in MATLAB incorporating the exact method. Effect size between groups was computed using Cohen’s *d* score, and the correlation coefficients between likelihood of seizure relapse and the severity of seizure outcome were determined using Spearman’s rank-order.

To enable reproducibility of our work, we will make available all the anonymised pre-surgery and surgically-spared brain networks of 51 patients, brain networks of 29 controls, codes for node abnormality computations, and all the trained machine learning models on the data presented in our manuscript (link to be generated upon acceptance of the manuscript).

## Results

The results are organised in three parts. First, we assessed if patients with greater number of abnormal nodes are predisposed to have suboptimal seizure outcome after surgery. Second, we investigated the effect of surgery in reducing the node abnormality load between the seizure-free and non-seizure groups. Third, we determined the generalisability of node abnormality measure, if it is to be incorporated in a clinical setting alongside other clinical attributes, to estimate the chances of seizure recurrence for new patients. The overall pipeline is shown in Figure 2 and explained in Methods. The full clinical data for each patient is provided in Table S1.

### Abnormality load corresponds with surgical outcome and post-surgery seizure relapse

We investigated the abnormality load in surgically-spared and pre-surgical networks. Figure 3(a-d) illustrates abnormal nodes in the surgically-spared networks for four patients. The patients in panels (a) and (b) were seizure free (ILAE 1) and having auras (ILAE 2) respectively at one-year after surgery and did not relapse subsequently; they had a relatively low node abnormality load. The patient in panel (c) initially had auras only (ILAE2) at one-year post-surgery but later relapsed; this patient showed a higher abnormal node count in the surgically-spared network. The patient in panel (d), with the highest number of abnormal nodes, had the worst surgical outcome of ILAE 5 at one year, which persisted upon follow-up. In these four cases a greater abnormality load was associated with seizure relapse, and worse outcomes.

**Figure 3:**
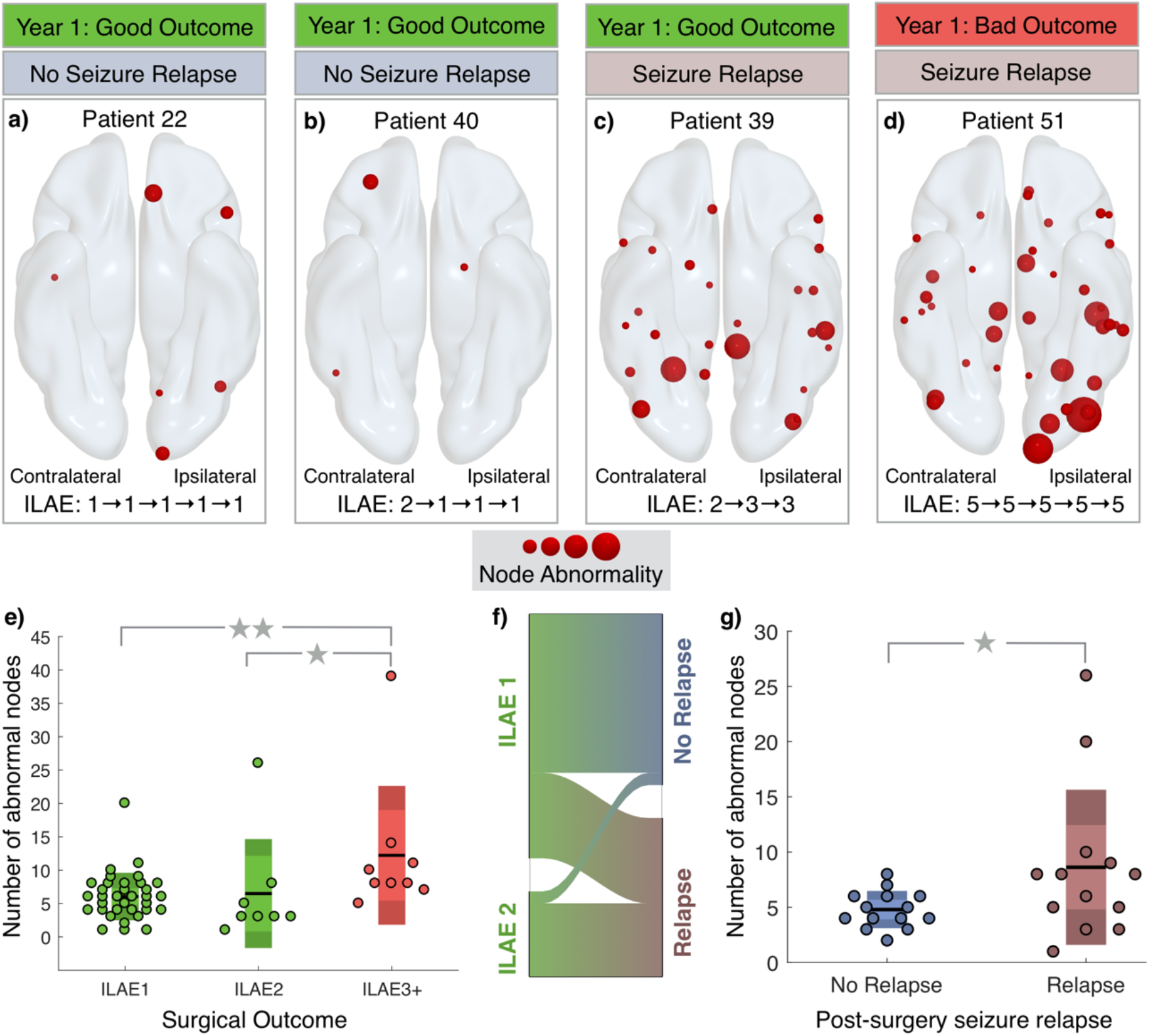
Association between the number of abnormal nodes in surgically-spared network with year-one surgical outcome and relapse. **a-d)** Four patients are shown with their year-one surgical outcome and relapse information. Panel **a)** and **b)** show fewer abnormal nodes in patients with ILAE 1 and ILAE 2 outcomes respectively with no relapse. Panel **c)** shows a patient with many abnormal nodes remaining yet having an ILAE 2 outcome at year-one but relapsing subsequently. Panel **d)** shows a large number of abnormal nodes remaining in a patient who was never seizure-free in five years. **e)** Significantly more abnormal nodes remained in ILAE 3+ patients compared to ILAE 1 and ILAE 2. **f)** Alluvial flow diagram showing proportion of relapsed patients with ILAE 1 or ILAE 2 at year-one. **g)** In ILAE 1-2 patients, those who relapsed had significantly more abnormal nodes in the surgically-spared network.

Figure 3(e) shows the node abnormality load in surgically-spared network for the entire patient cohort. Patients who were not seizure free (ILAE 3-6) at one-year post surgery, had significantly more number of abnormal nodes than patients who were seizure free (*p* = 0.002, *d* = 0.8 between ILAE 1 and ILAE 3-6 and *p* = 0.009, *d* = 0.6 between ILAE 2 and ILAE 3-6). Here, we chose to analyse ILAE 2 as a separate group because clinical data (Table 1, S1) suggests that these patients, albeit free from disabling seizures at year-one, have a greater propensity to relapse in later years (Fairclough *et al.*, 2018). Studying only the subset of patients who were seizure-free (i.e. ILAE 1, 2) at 1 year (Figure 3f-g), patients who relapsed had more abnormal nodes than the patients who did not relapse (*p* = 0.04, *d* = 0.75). Therefore, node abnormality can discriminate the patients in which seizures continued or recurred after surgery from the patients who were free from disabling seizures.

Node abnormality in Figure 3, computed from the surgically-spared network, was defined as the nodes with at least 10% of abnormal (*z* > 2.8) connections. At this choice of thresholds, the discrimination (AUC) between the seizure free and non-seizure free group was the highest. Comparable results are found for other threshold values (Supplementary Figure S1), and with an alternative network parcellation (Supplementary Figure S4). Thus, the discriminatory ability of node abnormality measure is consistent across the choice of threshold or the choice of parcellation scheme.

We found similar results in the pre-surgery networks. ILAE 3-6 patients had significantly more abnormal nodes than ILAE 1 patients (*p* = 0.02). However, the size of this effect was less pronounced than in the surgically-spared networks, with relatively poorer discriminatory ability (Supplementary Figures S1, and S2). Therefore, our findings suggest that the surgically-spared network, which is the surgically informed subnetwork of the pre-surgery network, is more discriminatory in identifying seizure free from non-seizure free patients.

### Surgery-related effect on reducing abnormality load

How much effect does surgery have on reducing the abnormality load? We investigated the *differences* between the surgically-spared, and pre-surgery networks in terms of their abnormality load, and whether the projected change in abnormality load due to surgery was greater and more widespread in one outcome group compared to another. The proportion of abnormal nodes in different brain areas for ILAE 1 (seizure free) and ILAE 3-6 (not seizure free) groups are shown in Figure 4.

**Figure 4:**
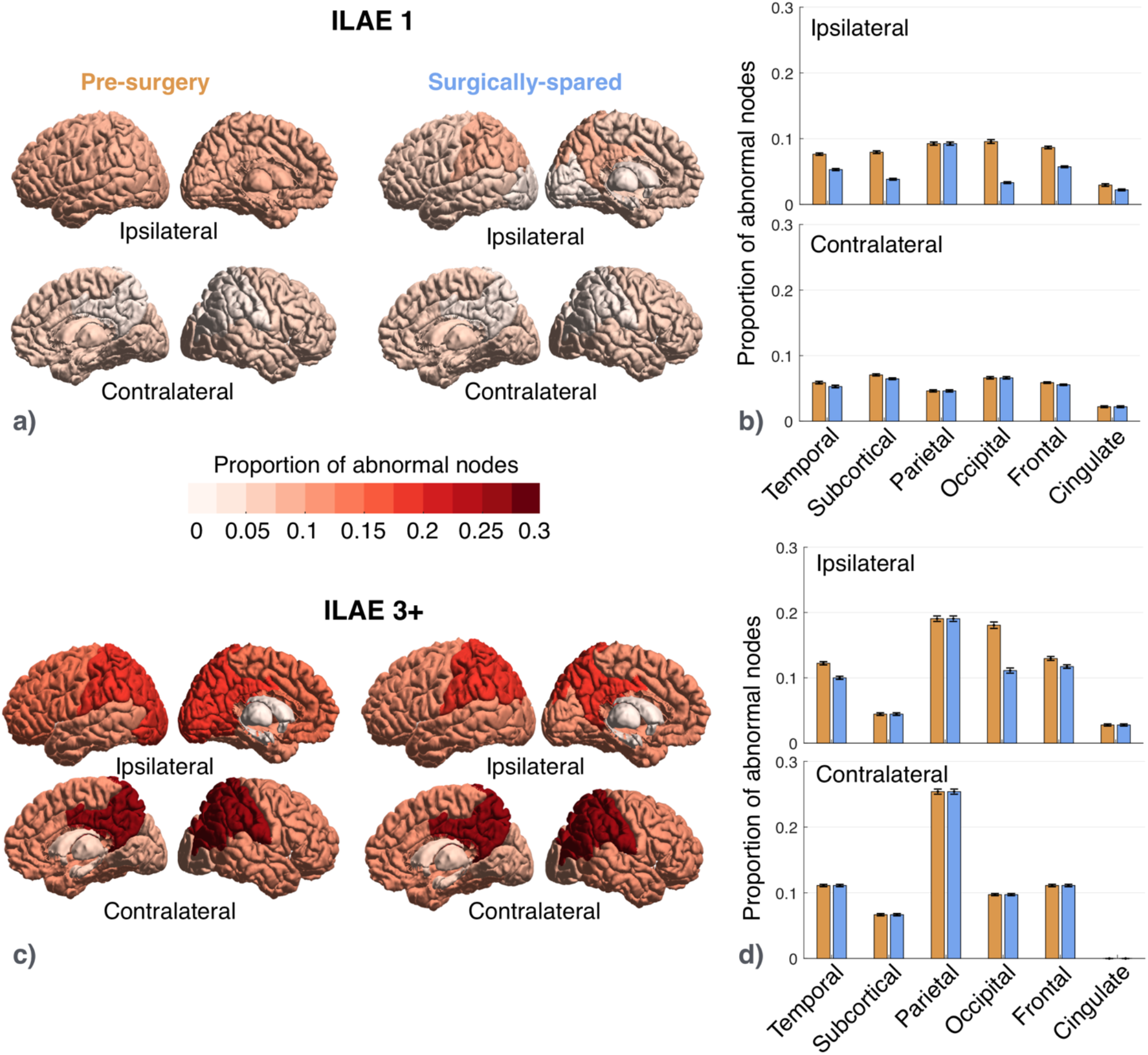
Effect of surgery in reducing node abnormality is more widespread in the seizure free group. **a)** Proportion of abnormal nodes computed for pre-surgery and surgically-spared networks are colour coded for six ipsilateral and contralateral brain areas in the seizure free (ILAE 1) group. **b)** Bar plot shows the drop in surgically-spared network compared to pre-surgery network in five ipsilateral (temporal subcortical, occipital, frontal, and cingulate) and two contralateral (temporal and sub-cortical) areas in the seizure free (ILAE 1) group. Error bars represent the standard error of the proportion of abnormal nodes in each area. **c)** Different brain areas in pre-surgery and surgically-spared network are colour coded based on the proportion of abnormal nodes in the non-seizure-free group (ILAE 3+) **d)** Corresponding bar plot showing a drop in node abnormality in surgically-spared network in three ipsilateral areas: temporal, occipital, and frontal in the not-seizure-free group (ILAE 3+).

In terms of the spatial extent of surgery, the expected reduction in the proportion of abnormal nodes was more widespread in the seizure free group than in the non-seizure free group. The ILAE 1 group had a drop in the proportion of abnormal nodes in the surgically-spared network, compared to the pre-surgical network, in seven areas: five ipsilateral and two contralateral (Figure 4a-b). In ILAE 3-6 group, however, the drop in the proportion of abnormal nodes was limited to three ipsilateral areas: temporal, occipital, and frontal (Figure 4c-d). Similar surgery related effect was found for node abnormality computed at different threshold values (Supplementary Figure S3).

In terms of reduction in the amount of abnormality load, ILAE 1 patients had larger proportional reductions than ILAE3-6 patients (*p* = 0.01), however, their absolute reduction did not differ significantly (*p* = 0.28) (Supplementary Figure S5). Thus, we suggest that the temporal lobe epilepsy surgery causes a greater and widespread reduction in abnormality load in the seizure free group than in the non-seizure free group.

### Personalised prediction of unsuccessful surgeries

We assessed the generalisability of the abnormality measure when used alongside other clinical attributes to predict patient-specific chances of poorer outcomes. Implementing nested-cross validation, we built machine learning models which classified new unseen (test) patients as either belonging to ILAE 1 or ILAE 3-6 group at 12 months. The model also scored each patient with a probability of belonging to either of the classes. Notably, the models were blind to three aspects of the data (a) all ILAE 2 patients, (b) ILAE classification 3, 4, 5 (the model simply sees these as ‘poor outcome’), and (c) outcomes at later years.

We incorporated up to 15 features in the model: 13 clinical attributes, the pre-surgical abnormality load, and the surgically-spared abnormality load. These features describe the presurgical attributes of patients and we evaluated them based on their combined ability in accurately predicting surgical outcomes at one-year. However, some features may be less informative than others in predicting surgical outcomes; including less informative features causes a drop in the prediction performance. Therefore, by implementing stepwise removal of less informative features, we obtained combinations of preoperative features that identified patients with poor seizure outcome at one-year after surgery in 100% cases (i.e., specificity). The area under the ROC curve (AUC) at every step of feature elimination is plotted in Figure 5(a) and magnified at one example point (marked with a star) in Figure 5(b) with the corresponding confusion matrix shown in the inset. Average prediction performance across all stepwise feature removals was robust at; AUC = 0.84 ± 0.07, accuracy = 0.79 ± 0.05, specificity = 0.89 ± 0.09, sensitivity = 0.77 ± 0.06. Supplementary Table 2 tabulates these prediction metrics in classifying seizure free and non-seizure free outcomes at every step. The lower panel in Figure 5(a) maps feature importance after each iteration of feature removal. The node abnormality in the surgically-spared network stood out as the most informative feature; it was more than 1.5 standard deviations away from the next most important features: age at surgery, and number of AEDs taken before surgery. Thus, including the abnormality measures to characterise pre-surgical attributes of intractable TLE patients led to a high and robust classification performance in predicting surgical outcomes at one-year after surgery.

**Figure 5:**
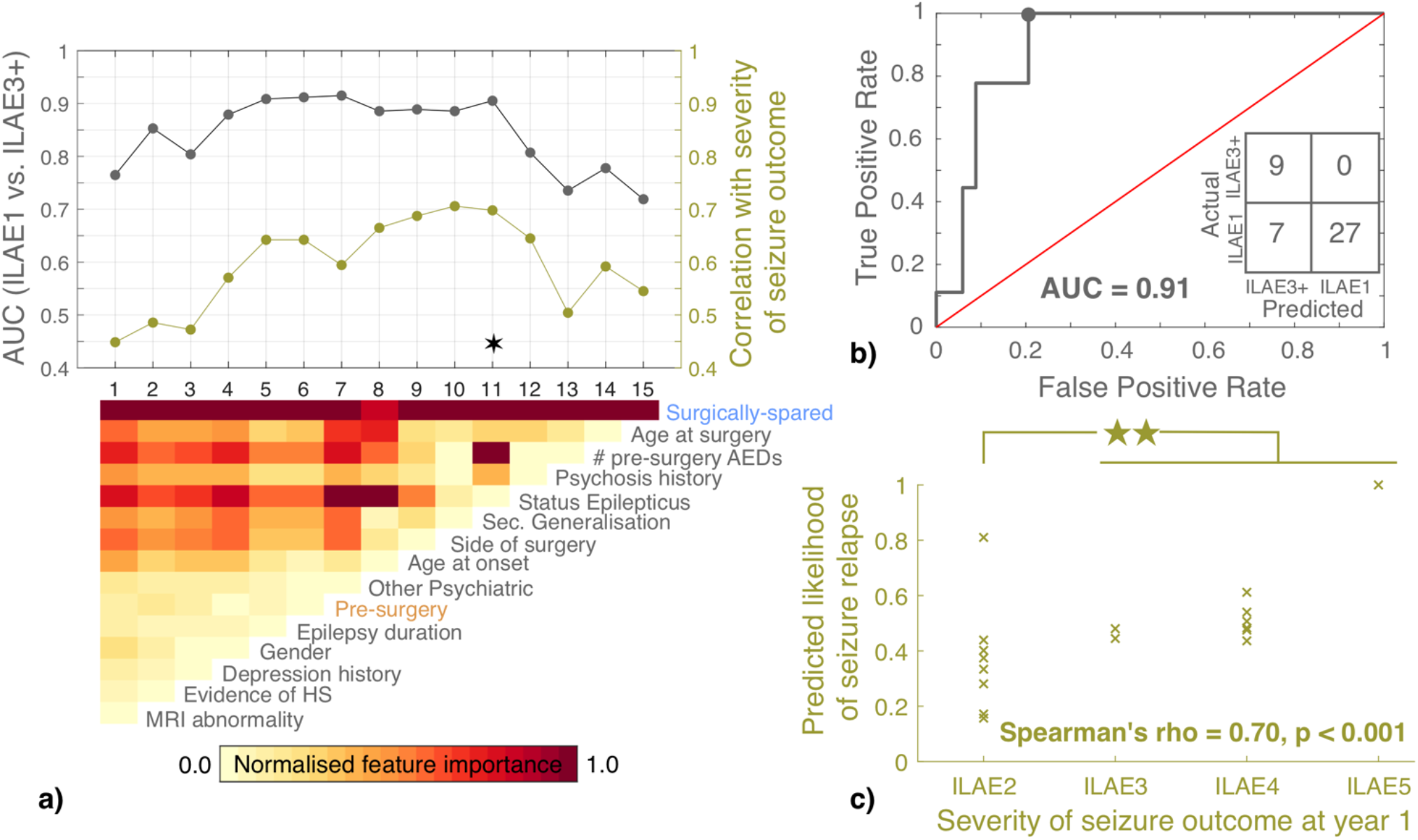
Prediction of seizure outcomes at year-one. **a)** AUC of SVMs that predicted seizure free (ILAE 1) and non-seizure free (ILAE3+) outcomes at one year after surgery are plotted in black. Blinded to the exact ILAE categories, the model predicted 12-month likelihood of seizure relapse for each patient. The Spearman’s rank correlation between likelihood of seizure relapse and the severity of surgical outcomes (ILAE class) at year-one are plotted in green. The lower panel of **a)** shows the relative feature importance of each SVM on a normalised scale between 0 and 1. The leftmost SVM, plotted at x = 1, incorporated all 15 features (13 clinical, node abnormality in pre-surgery and surgically-spared networks) to predict seizure free (ILAE 1) and non-seizure free (ILAE3+) outcomes at one year after surgery. Amongst all features, the relative importance of surgically-spared node abnormality was the highest whereas the relative importance of MRI abnormality was the least. Therefore, in the next iteration at x = 2, a new SVM was retrained using the 14 features, after removing the MRI abnormality feature. This stepwise removal of metrics was continued until only a single metric (surgically-spared node abnormality) remained. **b)** ROC curve is plotted at an example combination of features that yielded highest classification performance (AUC=0.91, specificity=1, sensitivity=0.79, accuracy=0.84). **c)** At the same example point, the correlation between the predicted likelihood of seizure relapse and the severity of seizure outcome at year 1 is shown. The predicted likelihood of seizure relapse was significantly different between ILAE 2 and ILAE 3-5 patients combined (*p* = 0.003, *d* = 0.95).

We next analysed the scores/probabilities assigned by the model to each patient to have a non-seizure free surgical outcome. Larger probabilities indicated a greater predicted likelihood of postoperative seizure at year 1 (i.e., the ILAE3+ group). Since the model was trained only on binary ILAE1 and ILAE3-6 outcomes it was blind to the spectrum of ILAE class data. We found that despite being blinded to such information, the predicted likelihood of seizure relapse was positively correlated with ILAE surgical outcome scale at year-one (Figure 5c). This positive association is consistent, even for the model trained using only the surgically-spared node abnormality feature (Figure 5a). Spearman’s *rho* values are plotted at each step-wise removal of features in Figure 5(a) and magnified for an example point in Figure 5(c). To confirm this result, we applied robust regression to obtain the regression slope and tested the significance of the steepness of the regression slope using permutation test (1000 permutations, *p* = 0.004 in Supplementary Figure S6). Therefore, our result shows that the pre-surgical clinical profile of patients, when assessed along with the abnormality measures, can inform about the grade of seizure outcomes which a patient would expect after surgery.

How informative are the pre-surgical features in predicting seizure recurrences in the long-term? We analysed this by checking the association between the predicted likelihood of seizure relapse and the actual relapse data for patients who were seizure free (ILAE 1,2) at year one. Patients who were not seizure free at year one (ILAE 3-6) were not included in the relapse category. We found no association with seizure relapse when the pre-surgical features of patients were characterized using a combination of clinical and network abnormality measures (Supplementary Figure S7). However, significant association with relapse was present at year 3, 4, and 5 when the pre-surgical features of patients were characterized using only the abnormality load in surgically-spared network (Figure 6). Hence, we suggest that in determining the long-term seizure recurrences, pre-surgical clinical attributes are less informative than the measure of abnormality load expected to be present in a patient after surgery.

**Figure 6:**
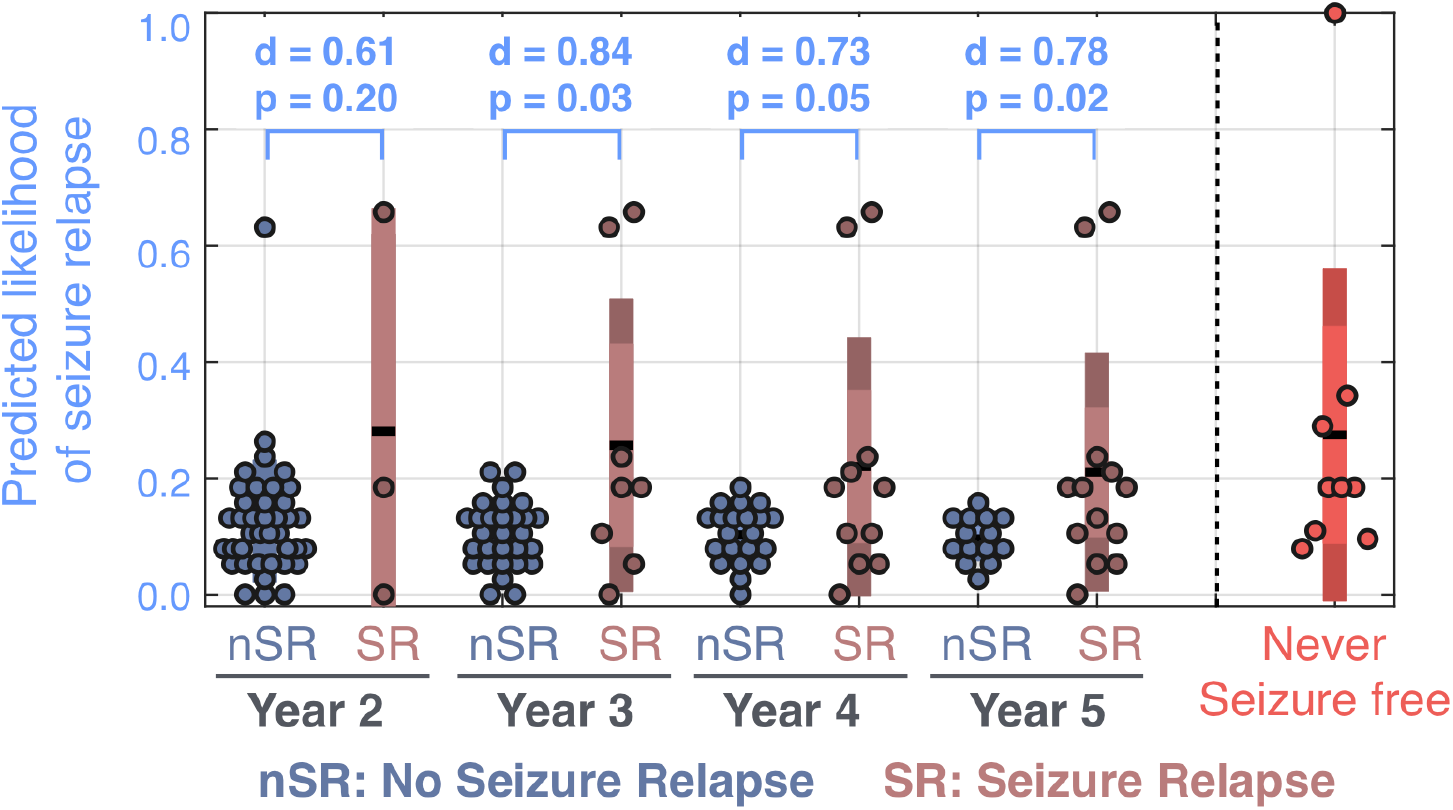
Predicted likelihood of seizure relapse at one-year was higher in patients who had seizure relapse at later years. The predicted 12-month likelihood of seizure relapse was estimated from the SVM model trained with only the surgically-spared node abnormality feature. The likelihood of seizure relapse for patients who were never seizure free (i.e. ILAE 3-6 at year 1) are shown in red. Amongst the patients who were initially seizure-free (i.e., ILAE 1 or ILAE 2 at year 1), higher likelihood of seizure relapse was predicted for those who had a subsequent relapse. This is despite the model being blinded to the outcomes at later years.

In summary, we achieved excellent performance in predicting seizure outcomes at one year when intractable TLE patients were assessed based on abnormality measures and clinical attributes. This combined pre-surgical profiling of patient attributes was also informative about the grades of seizure outcomes (ILAE class) at year one. However, beyond the first year after surgery, node abnormality in the surgically-spared networks were more informative about seizure relapses.

## Discussion

We investigated the association of surgical outcomes and relapse with the abnormality load computed in whole-brain pre-surgical network, and surgically spared subnetwork. Patients were more likely to have a poorer seizure outcome at one-year post surgery or a seizure relapse in five years, if more abnormal nodes were present in the surgically-spared network. Investigating the spatial effect of surgery on abnormality load, we found that the seizure free group of patients would have had a more widespread reduction of abnormal nodes. We found that the abnormality load in pre-surgery and surgically-spared networks, combined with clinical attributes of patients, generalised to predict 12-month postoperative seizure freedom with 100% specificity and 0.91 AUC. With this combined characterization of patient attributes, we predicted the likelihood of seizure relapse patient-specifically which were correlated with the ILAE class, hence, informative of the seizure outcome expected at 12 months after surgery. Finally, we showed that node abnormality located in the surgically-spared networks were particularly informative in identifying patients who were initially seizure free but would relapse after the first year of surgery and up to 5 years.

Altered white-matter tract integrity, extending beyond the ipsilateral temporal lobe to the extra-temporal and contralateral regions, has been extensively studied in TLE (Concha *et al.*, 2012; Otte *et al.*, 2012). Diffusion abnormalities with predictive value of postoperative outcomes are seen at the individual tract (Keller *et al.*, 2017), and also at the whole-brain network level (Bonilha *et al.*, 2013). It is indeed possible that areas which are anatomically normal, compared to controls, are epileptogenic via some other mechanism. Making an assumption that epileptogenic areas are most likely abnormal, our results show the presence of abnormal areas outside the temporal lobe which are spared by surgery. We suggest that net abnormality load, being a prognostic marker of seizure outcome, may be linked with epileptogenicity and have the potential to support epileptogenic networks.

A recent study on a different dataset with different imaging protocols, investigated network abnormality as a personalised predictor of surgical outcomes (Bonilha *et al.*, 2015). In that study, pre-surgery networks were constructed based on the number of streamlines connecting different regions. Similar to our study, connections between ROIs were normalised (z-transformed) against a control distribution. The similarity between our results suggest that: a) normalised patient networks using a local control distribution may enable reproducibility, comparison, and possibly grouping of patients between sites, and b) non-invasive personalised network biomarkers for predicting the likelihood of specific post-surgery outcomes in TLE are possible. We further showed the benefit of incorporating the information about the location of surgery to predict the surgical outcome.

The current standard for individualised prediction of surgical outcome primarily relies on clinical variables. However, there is considerable controversy in the literature regarding the presurgical clinical factors that may help predict surgical outcome. The review by (Bonilha and Keller, 2015) discussed discordant findings between different studies; features found predictive of seizure outcome in some studies are not predictive in others. Combining variables as nomograms, gave only modest concordance-statistics (c-statistics) of ∼ 0.5 on validation models (Jehi *et al.*, 2015). A more recent study estimated the probability of seizure freedom using combinations of up to 27 clinical variables on a mixed cohort of TLE and ETLE patients (Bell *et al.*, 2017). Our findings indicated that combining clinical variables with brain connectome derived features such as: abnormality load in pre-surgery and surgically-spared networks, can improve prediction of surgical outcomes in the short-term. Particularly for long-term predictions, the abnormalities in the surgically-spared networks, which are expected to remain after surgery, may be a more reliable measure because they associate with relapses. Hence, we propose to combine our node abnormality measure with clinical variables in a large mixed-cohort patient (Bell *et al.*, 2017) to improve estimation of the probability of seizure freedom/relapse after surgery.

In combining multivariate data, machine learning techniques delineate, rank, and fit input features of the training set to draw a decision boundary in a high dimensional space that maximises prediction (Bonilha *et al.*, 2015; Munsell *et al.*, 2015; Gleichgerrcht *et al.*, 2018; Taylor *et al.*, 2018; Morgan *et al.*, 2019). While a binary classification of seizure free and non-seizure free outcomes at 12 months is important, predicting long-term trajectories of seizure freedom is also crucial to inform clinical management decisions. In our study, the classifier not only predicted the surgical outcome at one-year but also predicted the likelihood of seizure relapse. This additional information may be clinically useful for advising patients about their chances of poor outcome post-surgery beyond the first 12 months and represents a key novelty of our work.

The outcome of epilepsy surgery will not just depend on the brain network before the surgery but also on the location and extent of surgery (Ji *et al.*, 2015; Taylor *et al.*, 2018). In this study, we retrospectively included the information of surgery by drawing a resection mask and inferring an expected surgically-spared network. A limitation of our work is that we are only inferring the *expected* postoperative network, rather than deriving it from postoperative dMRI data (Winston *et al.*, 2014). However, an analysis using *actual* postoperative dMRI data would only have very limited value in terms of improving the pre-operative decision making, since the outcome could only be seen after the surgery has been performed. In contrast, our approach can be used *before* the actual surgery to evaluate likelihood of success. A prospective application would involve drawing a resection mask for an *intended surgery* on sMRI of a patient acquired before surgery (Taylor *et al.*, 2018). Surgical data (either retrospectively delineated or prospectively planned) enables the study of expected changes after surgery (Taylor *et al.*, 2018). We showed that this information improves the prediction performance more so than just the pre-surgery networks which are naïve to surgical information. We envisage a software tool where multiple standard operations could be selected (e.g. selective amygdalohippocampectomy, or anterior temporal lobe resection) and their impact on the abnormality load compared. Tailored resections could also be tested to see what the effect of a larger resection might be. Such a tool could then be used to prospectively guide decision making regarding personalised resection strategies.

With regard to the extent of surgical resection, (Schramm, 2008) showed that the amount of tissue resected does not necessarily relate to improved surgical outcome. What is included in the resection, however, may have a significant influence on outcome (Siegel *et al.*, 1990; Bonilha *et al.*, 2012). The question arises: will a tailored resection, designed to reduce the number of abnormal nodes, lead to a better outcome? While more investigations are needed to confirm this hypothesis, we found that the seizure free patient group had a more widespread reduction of abnormality load due to surgery. Simulated computer models may facilitate a more detailed analysis to investigate alternative surgical strategies in a personalised manner (Sinha *et al.*, 2017).

Our findings must be interpreted in the context of some caveats. Node abnormality may be representative of a) network reorganisation in response to seizures, b) neurodegenerative process due to seizures, c) structures facilitating seizures, or combinations of (a)-(c). In our study we could not disentangle these aspects with respect to node abnormality. We did not detect any significant correlation between clinical variables and node abnormalities. Though our sample size is reasonably large (Bonilha *et al.*, 2015; Munsell *et al.*, 2015; Keller *et al.*, 2017; Gleichgerrcht *et al.*, 2018), it is not of the size of typical epidemiological studies. Neural architecture depends on several subject-specific factors including language dominance, handedness, and other physiological variables. These relationships may further influence the node abnormality measure. Thus, our results should motivate a larger study to test its generalisability, ideally across multiple sites. Finally, we highlight, based on the pre-surgical, surgically-spared networks, and clinical variables, the chances of *at least one* relapse in five-years. However, the trajectory of seizure remission and relapse is more complicated. Patients may have repeated remissions and relapses due to drug-effects, environmental factors, or other causes.

In summary, we have shown evidence of node abnormality being an important non-invasive marker of surgical outcome and its severity at year one post-surgery. Node abnormality is also related with likelihood of seizure relapse in long-term. We demonstrate improvement in prediction performance when including surgery information with the pre-surgery network and clinical data. We believe this to be an important step towards complementing clinical decision making on patient and surgery selection for intractable TLE as well as for patient counselling regarding the risks of seizure severity expected after surgery.

## Acknowledgements

NS was supported by Research Excellence Academy, Newcastle University, UK. PNT was supported by the Wellcome Trust (105617/Z/14/ Z and 210109/Z/18/Z). YW was supported by the Wellcome Trust (208940/Z/17/Z). We thank Gabrielle Schroeder, Pawel Widera, Sriharsha Ramaraju, other members of ICOS, and CNNP lab (www.cnnp-lab.com) for discussions. SBV was funded by the UCLH NIHR BRC. Scan acquisition and GPW were supported by the MRC (G0802012, MR/M00841X/1). We are grateful to the Epilepsy Society for supporting the Epilepsy Society MRI scanner. This work was supported by the National Institute for Health Research University College London Hospitals Biomedical Research Centre.

