## SupplementaryMaterial for "Node abnormality predicts seizure outcome and relates to long-term relapse after epilepsy surgery"

#### Supplementary Figures

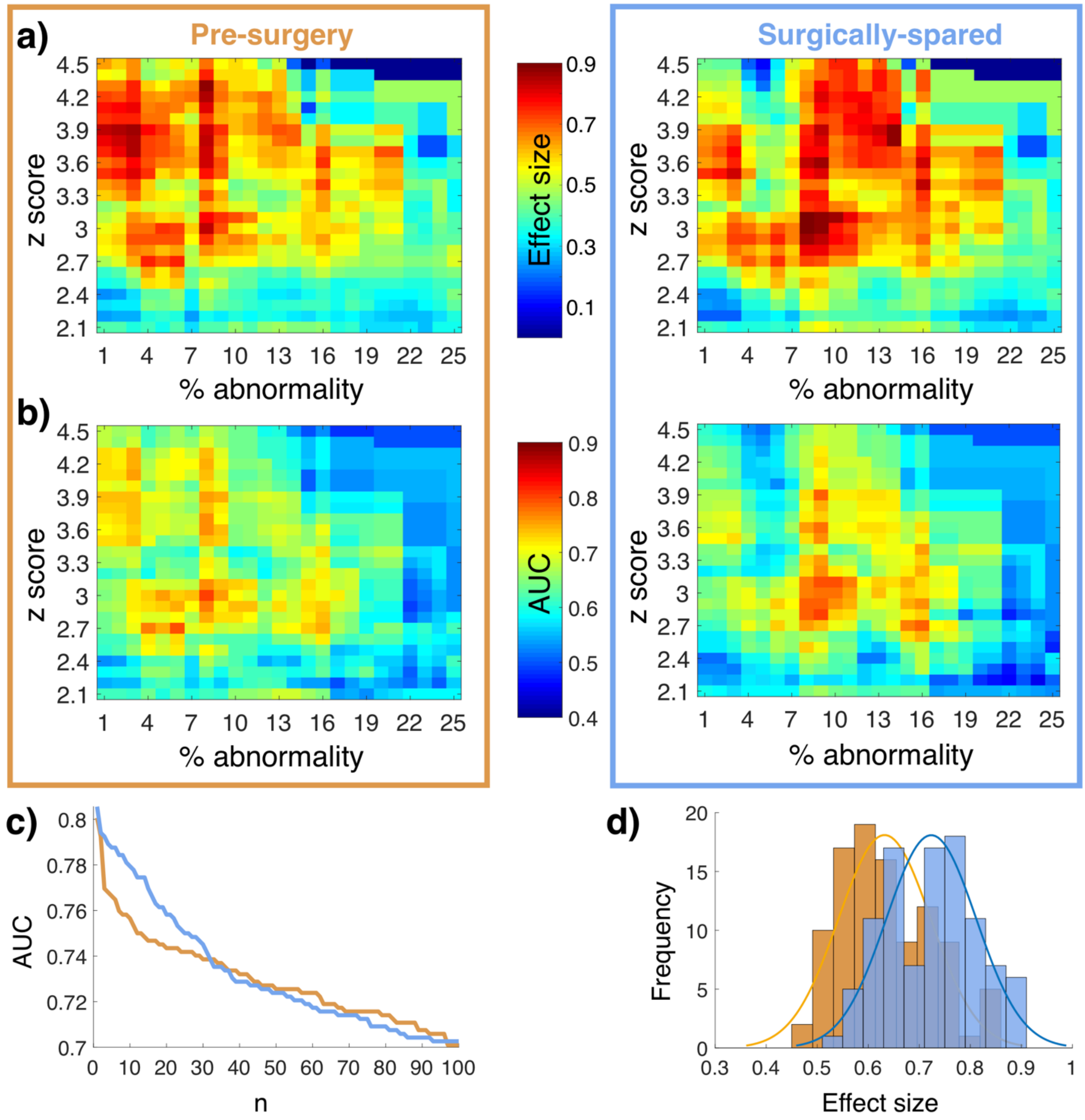


**Figure S1: Association between node abnormality and surgical outcomes in pre-surgery and surgically-spared network is consistent across thresholds. a)** Effect size (d-score) for node abnormality between ILAE 1 and ILAE 3+ groups in pre-surgery and surgically-spared networks. Positive effect size indicates higher node abnormality in ILAE 3+ patients compared to ILAE 1 patients. Medium to large positive effects size, colour coded in red, are evident across a large range of thresholds. **b)** AUC quantifying the discriminatory value of node abnormality is shown at every point on the threshold grid for pre-surgery and surgically-spared networks. **c)** 100 highest AUCs sampled from the threshold grid of pre-surgery and surgically-spared networks in panel **b)** are plotted against each other. Relatively higher AUCs are apparent in surgically-spared networks. **d)** Histogram of effect size sampled from pre-surgery and surgically-spared networks corresponding to the threshold values of 100 highest AUCs of surgically-spared network.

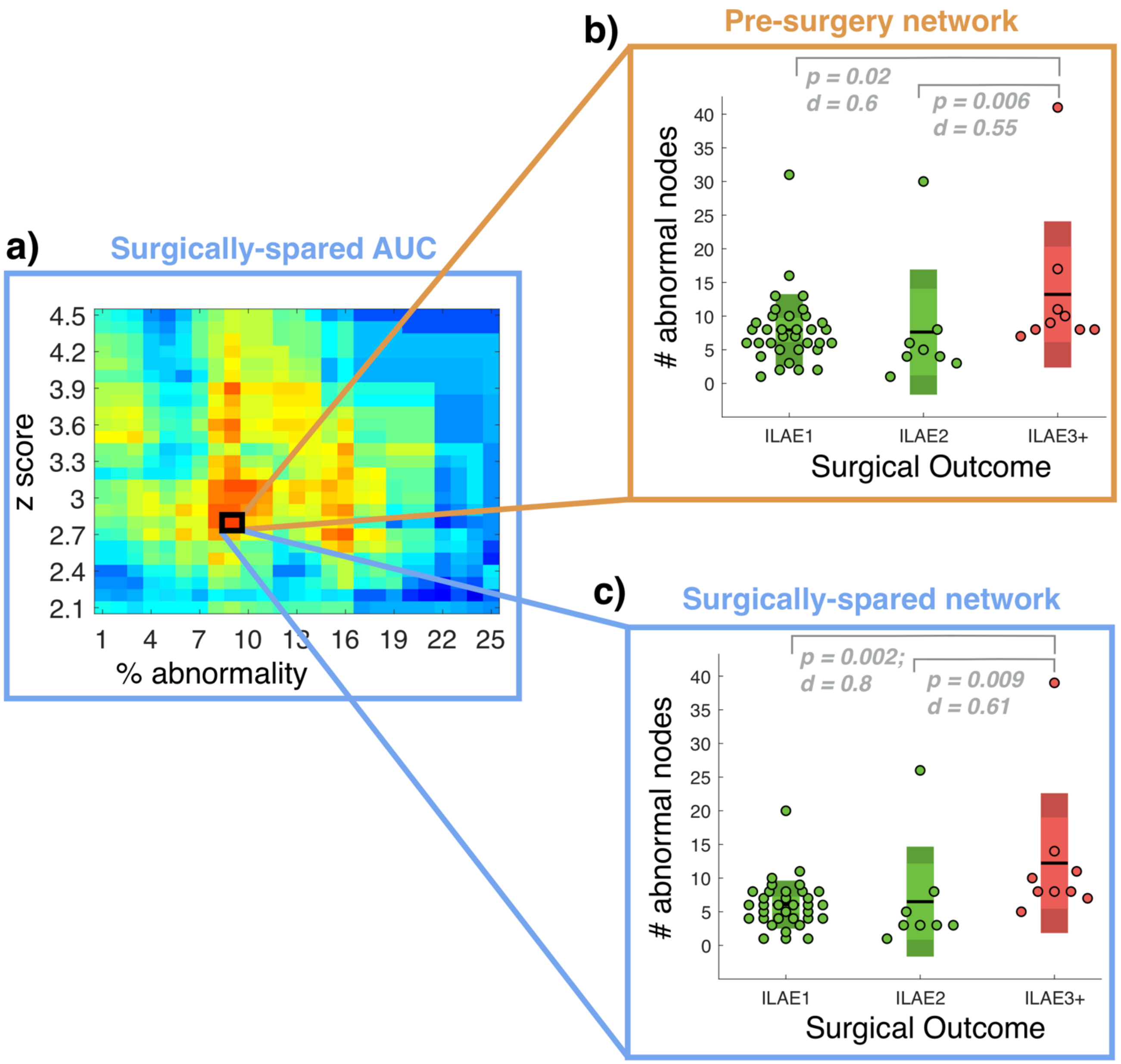


**Figure S2: Surgically-spared networks are more discriminatory than the pre-surgery networks.** **a)** Corresponding to the threshold of highest AUC, results are shown for pre-surgery and surgically-spared networks. **b)** Node abnormality computed from pre-surgery network discriminates ILAE1 from ILAE3+ with an effect size of d = 0.6 and ILAE 2 from ILAE 3+ with an effect size of d = 0.55. In comparison, node abnormality computed from surgically-spared networks in **c)** discriminate ILAE1 from ILAE3+ with an effect size of d = 0.8 and ILAE 2 from ILAE 3+ with an effect size of d = 0.61.

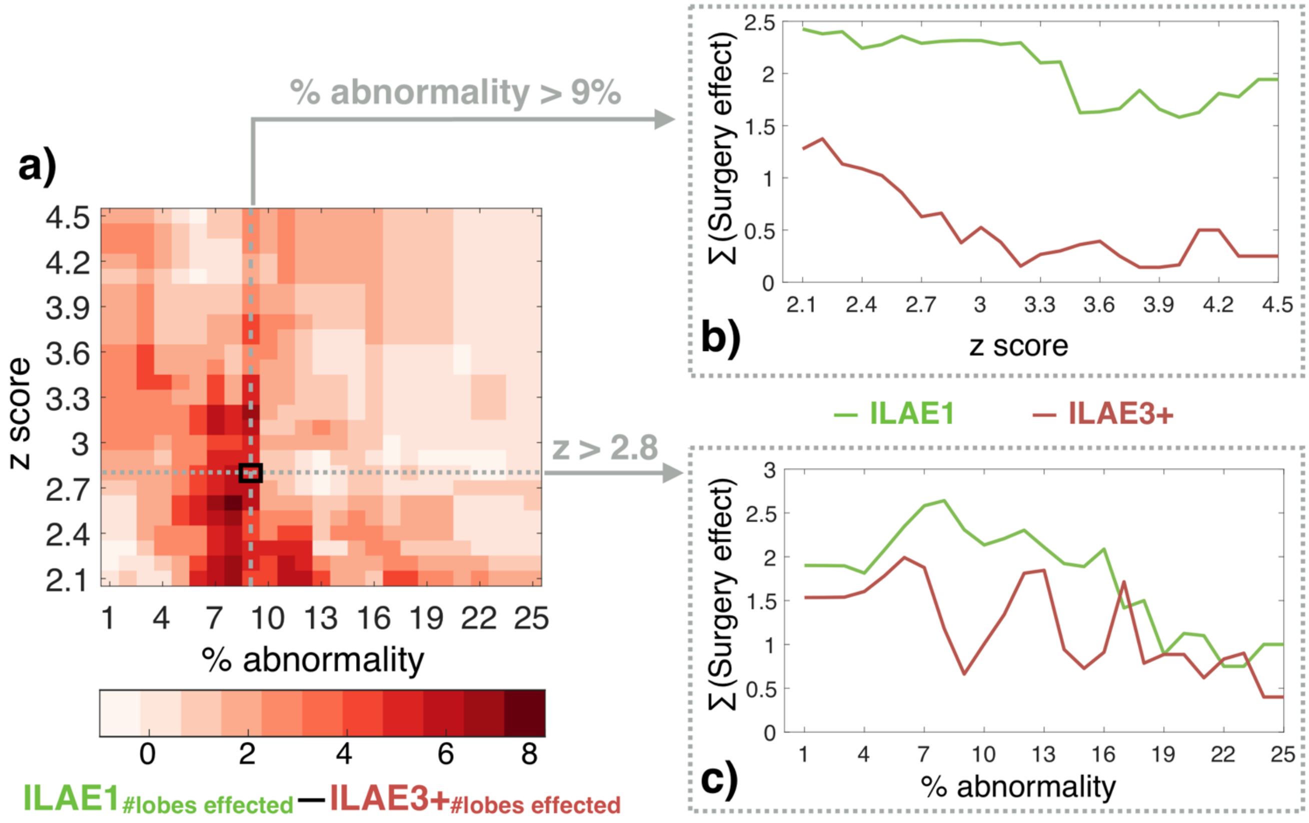


**Figure S3: The widespread effect of surgery in reducing node abnormality in seizure free group is consistent across the thresholds.** Furthering our results in Figure 4, we show the difference between the number of lobes effected in seizure free and non-seizure free group across thresholds in panel **a)**. Positive differences in **a)** indicate that surgery effects the node abnormalities in more brain areas of ILAE 1 group compared to ILAE 3+ group. Along the two slices taken from the grid, the net effect of surgery is shown between ILAE 1 and ILAE 3+ groups in panel **b)** and **c)**. The effect of surgery at each lobe is computed as: 1 – ratio of proportion of abnormal nodes in ILAE1 to ILAE3+. The net effect of surgery is computed as the sum of the aforementioned ratio across all the ipsilateral and contralateral brain areas.

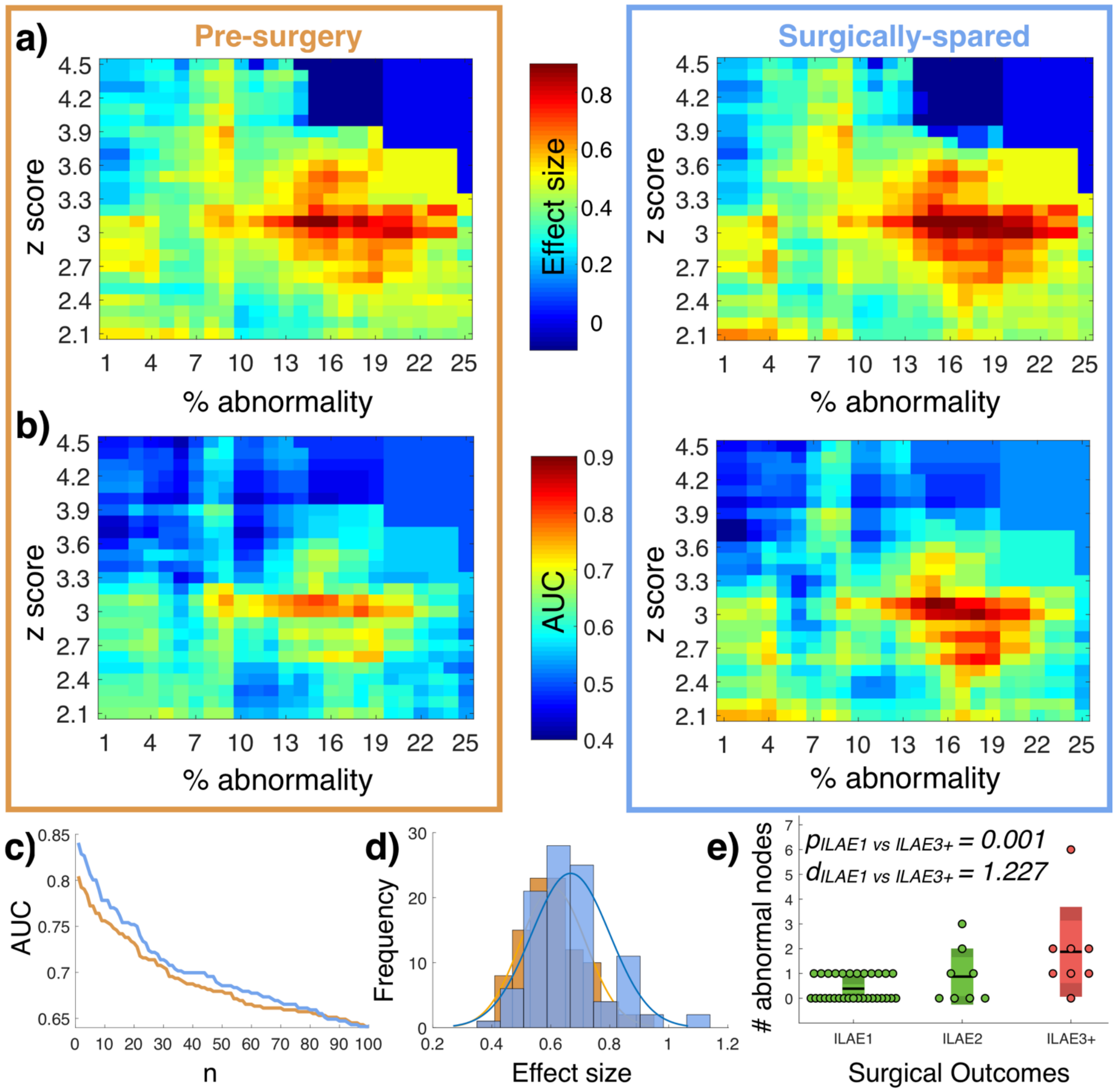


**Figure S4: With networks inferred using Desikan-Killiany parcellation scheme the association between node abnormality and surgical outcomes in pre-surgery and surgically-spared networks are consistent across thresholds. a)** Effect size (d-score) for node abnormality between ILAE 1 and ILAE 3+ groups in pre-surgery and surgically-spared networks. Positive effect size indicates higher node abnormality in ILAE 3+ patients compared to ILAE 1 patients. Medium to large positive effects size, colour coded in red, are evident across a large range of thresholds. **b)** AUC quantifying the discriminatory value of node abnormality is shown at every point on the threshold grid for pre-surgery and surgically-spared networks. **c)** 100 highest AUCs sampled from the threshold grid of pre-surgery and surgically-spared networks in panel **b)** are plotted against each other. Relatively higher AUCs are apparent in surgically-spared networks. **d)** Histogram of effect size sampled from pre-surgery and surgically-spared networks corresponding to the threshold values of 100 highest AUCs of surgically-spared network. **e)** At an example point on the threshold grid corresponding to the highest AUC, the box plot shows significantly higher number of abnormal nodes in ILAE 3+ patient than in ILAE 1 patients.

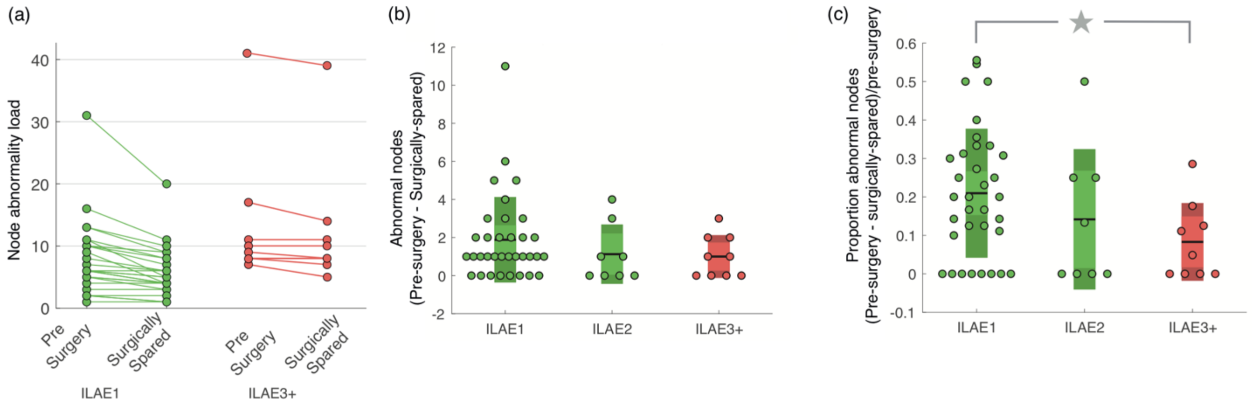


**Figure S5: Change in node abnormality load between pre-surgery and surgically-spared networks.** a) Reduction in the number of abnormal nodes between pre-surgery and surgically-spared networks are shown for ILAE 1 and ILAE 3+ patients. ILAE 3+ patients have a greater number of abnormal nodes than ILAE 1 patients. Inspecting visually, the reduction of abnormality load, apparent by the slope of the lines, are more in ILAE 1 patients than in ILAE 3+. b) The absolute reduction in abnormal nodes between pre-surgery and surgically spared networks are higher on average in ILAE 1 patients but not statistically significant ($p = 0.28$). c) The drop in the number of abnormal nodes between pre-surgery and surgically-spared networks relative to the node abnormality load pre-surgery is significantly higher in the ILAE 1 patients compared to the ILAE 3+ patients ($p = 0.01$).

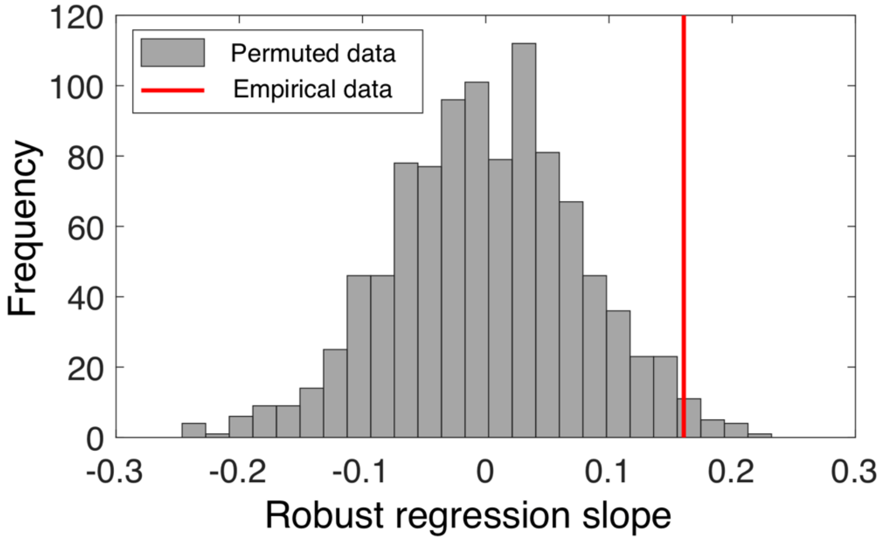


**Figure S6:** Steepness of regression slope obtained from robust regression tested for significance using permutation testing (p = 0.004, number of permutations = 1000).

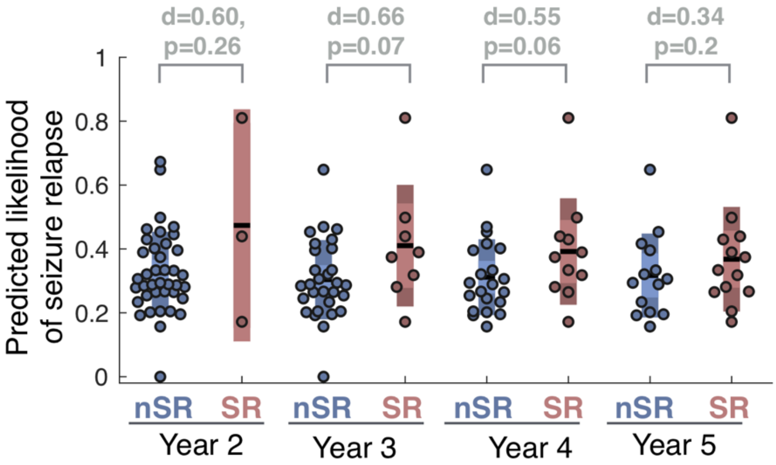


**Figure S7: On combining network features with clinical attributes association with relapse is lost.** The predicted 12-month likelihood of seizure relapse was estimated from the SVM model at an example combination of features that yielded highest classification performance (corresponding to Figure 5b-c). Amongst the patients who were initially seizure-free (i.e., ILAE 1 or ILAE 2 at year 1), the likelihood of seizure relapse was not different to those who had a subsequent relapse.

### Supplementary Methods

#### Imaging protocols

MRI data were acquired on a 3T GE Signa HDx scanner (General Electric, Waukesha, Milwaukee, WI). Standard imaging gradients with a maximum strength of $40\,mTm^{-1}$ and slew rate $150\,Tm^{-1}s^{-1}$ were used. All data were acquired using a body coil for transmission, and 8-channel phased array coil for reception. Standard clinical sequences were performed including a coronal 3D T1-weighted volumetric acquisition (matrix, $256 \times 256 \times 170$; in-plane resolution, $0.9375 \times 0.9375\,mm$ with slice thickness 1.1 mm).

Diffusion MRI data were acquired using a cardiac-triggered single-shot spin-echo planar imaging sequence (Wheeler-Kingshott et al., 2002) with echo time $= 73\,ms$. Sets of 60 contiguous $2.4\,mm$-thick axial slices were obtained covering the whole brain, with diffusion sensitizing gradients applied in each of 52 noncollinear directions (b-value of $1200\,mm^2s^{-1}$ $[\delta = 21ms, \Delta = 29\,ms$ using full gradient strength of $40\,mTm^{-1}]$) along with 6 non-diffusion weighted scans. The gradient directions were calculated and ordered as described elsewhere (Cook et al., 2007). The field of view was $24 \times 24\,cm$, and the acquisition matrix size was $96 \times 96$, zero filled to $128 \times 128$ during reconstruction, giving a reconstructed voxel size of $1.875 \times 1.875 \times 2.4\,mm$. The DTI acquisition time for a total of 3480 image slices was approximately 25 min (depending on subject heart rate). These protocols are identical to our previous study (Taylor et al. 2018).

#### Pre-processing pipeline

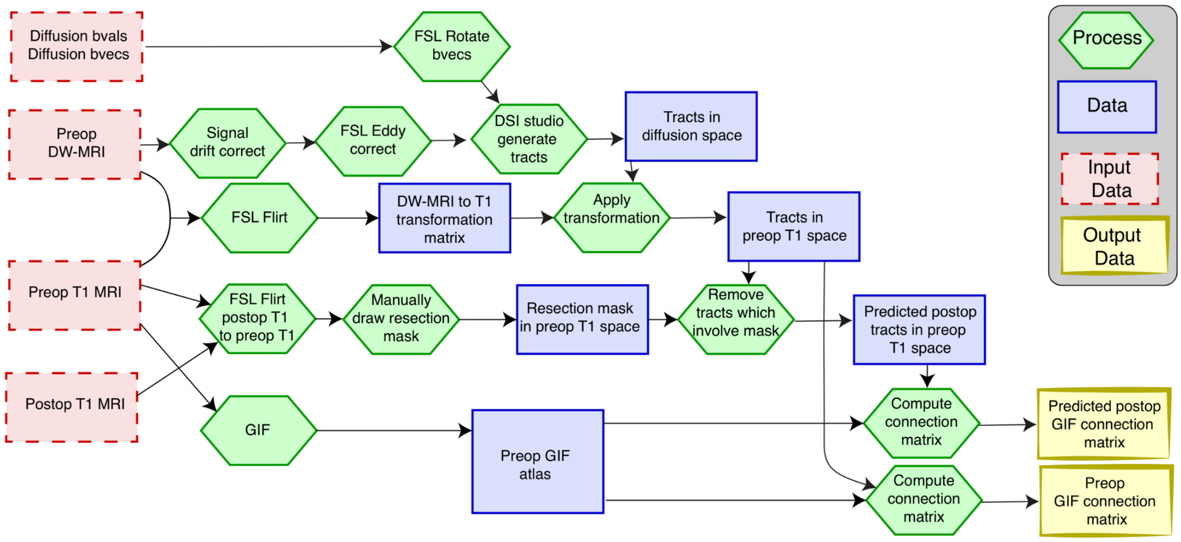


*Reproduced from Taylor et al. 2018:* Summary of processing pipeline to generate GIF network. The pipeline is applied to each subject.

We applied the same image pre-processing steps that we previously established on this data in our recent study (Taylor et al. 2018). The image processing pipeline is summarised in the flowchart reproduced from Taylor et al. 2018.

Preoperative diffusion MRI data were first corrected for signal drift (Vos *et al.* 2017), then eddy current and movement artefacts were corrected using the FSL eddy_correct tool (Andersson and Sotiropoulos 2016). The b vectors were then rotated appropriately using the 'fdt-rotate-bvecs' tool as part of FSL (Jenkinson *et al.* 2012, Leemans & Jones, 2009). The diffusion data were reconstructed using generalised q-sampling imaging (Yeh, Wedeen, and Tseng 2010) with a diffusion sampling length ratio of 1.2. A deterministic fibre tracking algorithm (Yeh et al. 2013) was then used, allowing for crossing fibres within voxels, with seeds placed at the whole brain. Default tractography parameters from the 14 February 2017 build of DSI studio software were used as follows. The angular threshold used was 60 degrees and the step size was set to 0.9375mm. The anisotropy threshold was determined automatically by DSI Studio. Tracks with length less than 10 mm and more than 300mm were discarded. A total of 1,000,000 tracts were calculated per subject and saved in diffusion space.

To align the tracts with the ROIs we linearly registered the first non-diffusion-weighted image to the preoperative T1 image and saved the inverse of this transformation matrix using FSL FLIRT. We then multiplied every coordinate in every tract by this transformation matrix to get the tracts in T1 space. The quality of the registration between tracts, ROIs, and the resection mask was confirmed through visual inspection for all subjects. Since networks are constructed in native space, this removes any mismatching of track types due to potential nonlinear registration issues which is advantageous compared to previous studies of network change (Kuceyeski 2016). To generate preoperative connectivity matrices, we looped through all tracts and deemed two regions as connected if the two endpoints of the tract terminate in those regions. This generated a weighted connectivity matrix in which each entry in the matrix represents the number of streamlines connecting two regions. To generate predicted postoperative connectivity matrices, we performed the same process as above with one exception. Any tract that had any point within the resection mask was excluded from building the matrix. The inferred postoperative network therefore always had fewer streamlines than the preoperative network.

#### Inferring imprecise AED information

For one patient, the clinical metadata was incomplete—the number of AEDs taken by patient 43 was imprecise. To infer a value for the number of AEDs for this patient, we investigated the predictive power of the remaining clinical data for AEDs prediction using multiple linear regression. The fitting of the linear regression model was achieved using stats package in R, which uses the Least Squares method to minimise the sum of the squares of residuals. Initially, we included all available clinical data in the linear regression model. However, to overcome multicollinearity and remove redundant predictors we discarded, at each round, the predictor with highest *p*-value, until the regression model and its coefficients were statistically significant. A significant model (p = 0.002) with normally distributed residuals (Shapiro-Test: p = 0.7159) was obtained using epilepsy duration as single significant predictor. The model was used to predict, for the patient with missing data, the number of AEDs and its 95% confidence interval.

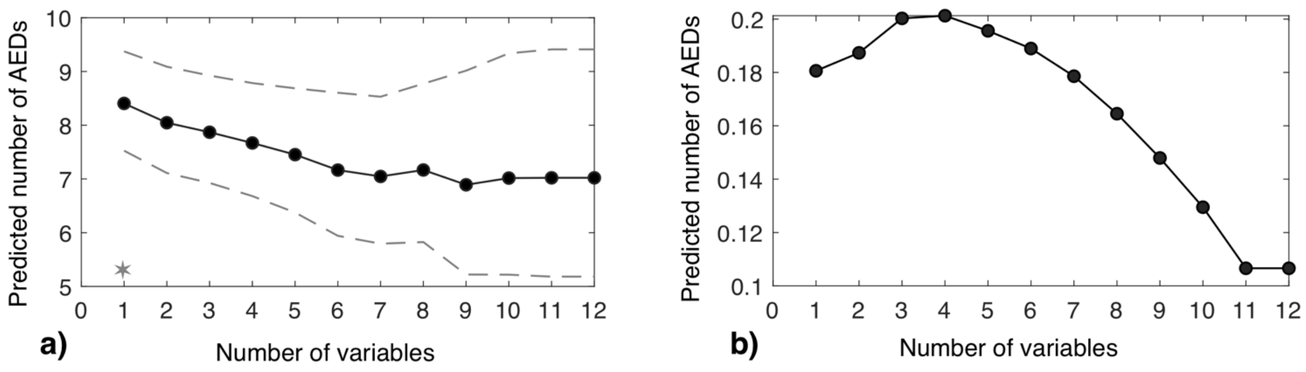


**Figure 1.** Prediction of number of AEDs for the patient with imprecise AED data. The number of variables represents the number of predictors used by the model at each round. In panel **a)** the solid line represents the predictive AEDs value and the dash lines the upper and lower confidence interval. The adjusted $r^2$ of the model at each round is shown in panel **b)**.

Since number of AEDs cannot be a fraction, we predicted the number of AEDs taken by patient 43 was 8. The following figure shows that the prediction performance remained similar regardless the imprecise data being imputed, left unchanged, or with patient 43 removed from the analysis. Thus, our results are robust to this missing data.

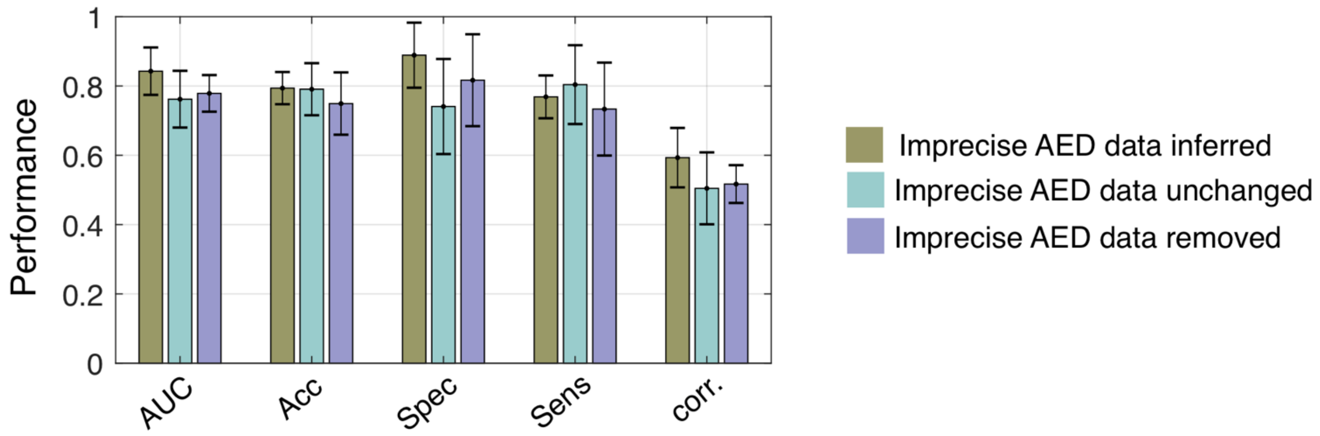


**Figure 2:** Consistent prediction performance was achieved regardless of how the imprecise AED data for patient 43 was treated. The bar plot shows the average performance metrics of SVMs in terms of AUC, accuracy, specificity, and sensitivity. The error bars represent the standard deviation across each step-wise feature removal.

#### Supplementary Tables

| Keywords | Definition |
|---|---|
| **Pre-surgery network** | Whole-brain connectivity using pre-operative diffusion MRI and pre-operative structural MRI only (Bonilha et al. 2015) |
| **Surgically-spared network** | Whole-brain connectivity using pre-operative diffusion MRI, pre-operative structural MRI, and a resection mask drawn on pre-operative structural MRI highlighting the surgery location (Taylor et al., 2017) |
| **Edge abnormality** | In the whole-brain connectivity the deviation of edges from normality. Normal distribution for the edges is inferred from a control population and deviation is measured in terms of z-score (Bonilha et al. 2015). |
| **Node abnormality** | Number of abnormal edges of a node normalised by the total number of edges in that node. |
| **Abnormality load** | The total number of abnormal nodes in the whole-brain connectivity of a patient. |

**Table S1: Complete preoperative clinical information of all patients with postoperative ILAE seizure outcomes and relapse**

[Attached separately]

**Table S2: Summarising SVM performance estimates at each step-wise feature removal**

| Feature elimination step | AUC | Accuracy | Sensitivity | Specificity | Correlation year-1 severity | |
|---|---|---|---|---|---|---|
| | | | | | Spearman's rho | p |
| 1 | 0.76 | 0.67 | 0.62 | 0.89 | 0.45 | 0.0355 |
| 2 | 0.85 | 0.79 | 0.76 | 0.89 | 0.49 | 0.0241 |
| 3 | 0.80 | 0.79 | 0.76 | 0.89 | 0.47 | 0.0278 |
| 4 | 0.88 | 0.86 | 0.85 | 0.89 | 0.57 | 0.0084 |
| 5 | 0.91 | 0.84 | 0.82 | 0.89 | 0.64 | 0.0027 |
| 6 | 0.91 | 0.84 | 0.82 | 0.89 | 0.64 | 0.0027 |
| 7 | 0.92 | 0.79 | 0.74 | 1.00 | 0.59 | 0.0059 |
| 8 | 0.89 | 0.79 | 0.74 | 1.00 | 0.66 | 0.0018 |
| 9 | 0.89 | 0.81 | 0.79 | 0.89 | 0.69 | 0.0011 |
| 10 | 0.89 | 0.79 | 0.74 | 1.00 | 0.71 | 0.0008 |
| 11 | 0.91 | 0.84 | 0.79 | 1.00 | 0.70 | 0.0009 |
| 12 | 0.81 | 0.79 | 0.79 | 0.78 | 0.64 | 0.0026 |
| 13 | 0.74 | 0.74 | 0.74 | 0.78 | 0.50 | 0.0195 |
| 14 | 0.78 | 0.74 | 0.71 | 0.89 | 0.59 | 0.0062 |
| 15 | 0.72 | 0.81 | 0.85 | 0.67 | 0.55 | 0.0118 |
